# FZD2 inhibits YAP and prevents cell cycle reentry in adult murine cardiomyocytes

**DOI:** 10.1101/2024.07.26.605158

**Authors:** Joseph A. Bisson, Pearl J. Quijada, Yves T. Wang, Janet K. Lighthouse, Jay Christian Helt, Eugene Kim, Edward E. Morrisey, Paul S. Brookes, Eric M. Small, Ethan D. Cohen

**Affiliations:** Department of Medicine, University of Rochester Medical Center, Rochester, NY 14642, USA; Department of Pharmacology and Physiology, University of Rochester Medical Center, Rochester, NY 14642, USA; Department of Surgery, Weill Cornell Medicine, New York, NY, 10065, USA; Aab Cardiovascular Research Institute, University of Rochester Medical Center, Rochester, NY 14642; Integrative Biology and Physiology, Division of Life Sciences, University of California, Los Angeles, 90095, USA; Department of Pharmaceutical Sciences, Wegmans School of Pharmacy, St. John Fisher College, Rochester, NY 14618, USA; Division of Digestive and Liver Diseases, Department of Medicine, Columbia University Medical Center, New York, NY 10032, USA; Department of Medicine and Cell and Developmental Biology, University of Pennsylvania, Philadelphia, PA 19104, USA; Department of Pediatrics, University of Rochester Medical Center, Rochester, NY 14642, USA

**Keywords:** Basic Science Research, Cell Signaling/Signal Transduction, Myocardial Regeneration, Myocardial Infarction, Genetically Altered and Transgenic Models

## Abstract

**Rationale:** Fully differentiated cardiomyocytes (CMs) are post-mitotic and cannot repopulate damaged tissue after myocardial infarction (MI). Understanding the mechanisms preventing CM proliferation or promoting their survival after injury may lead to treatment strategies for MI. While the effects of canonical WNT/β-catenin signaling in adult CMs have been examined, roles for non-canonical WNT signaling in cardiac homeostasis and repair remain unexplored.

**Objective:** To determine the function of the non-canonical WNT receptor frizzled 2 (FZD2) in adult cardiac homeostasis and injury.

**Methods and Results:** FZD2 was deleted from the CMs of adult mice to investigate its role in myocardial homeostasis. *Fzd2* conditional knockout (CKO) mice had cardiomegaly but not hypertrophy. FZD2-deficient CMs expressed proliferation and cytokinesis markers, suggesting that they have increased proliferation potential. FZD2-deletion caused the accumulation of β-catenin. However, β-catenin localized to the membranes of FZD2-deficient CMs and did not activate target gene expression. Instead, the YES-associated protein (YAP) regulated genes *v-myc avian myelocytomatosis viral oncogene 1 (Mycl)*, and *B cell leukemia/lymphoma 2 (Bcl2l1)* were upregulated in *Fzd2* CKO CMs relative to controls. Knockdown of FZD2 increased YAP activity in neonatal ventricular CMs (NVCMs), while overexpressing FZD2 inhibited YAP. Neither β-catenin knockdown nor mutating the large tumor suppressor 1 and 2 (LATS1/2) target site on YAP blocked the effects of FZD2 on YAP in NVCMs, suggesting that FZD2 utilizes different effectors than canonical WNT and Hippo signaling. *Fzd2* CKO and control mice were subjected to MI to determine if FZD2-deletion affects cardiac repair. While ischemia and necrosis were similar 24 hours post MI, *Fzd2* CKO mice had better cardiac function and less scarring than controls.

**Conclusions:** FZD2 reduces YAP activity and prevents adult murine CMs from reentering the cell cycle. FZD2-deletion improves heart function and reduces scarring in mice after MI, implicating FZD2 as a target for pharmacological intervention.

## INTRODUCTION

Heart disease is the leading cause of death in the industrialized world, killing one in four adults^1,2^. The most common form of heart disease is coronary artery disease, which can lead to myocardial infarction (MI) and the subsequent death of cardiomyocytes (CMs) due to ischemia^3^. CMs of the adult mammalian heart are terminally differentiated cells that are permanently lost to injury, and while most MI patients survive the initial ischemic event, many later succumb to heart failure^4^. Treatment options currently include drugs and mechanical devices that improve the heart’s function in a damaged state but do not involve preventing CM cell death or restoring those lost to ischemia. Improving our understanding of the molecular pathways that can be safely manipulated to protect against CM loss during MI would have a great impact on advancing therapies for patients.

In contrast to adult CMs, newborn mammalian CMs are proliferative and will regenerate damaged myocardium^5–9^. This capacity of regeneration and CM proliferation present in the neonatal heart is controlled by various developmental pathways that are not fully understood. One such pathway, the Hippo pathway, mediates the contact-dependent inhibition of cell proliferation and restricts organ size^10–12^. In Hippo signaling, mammalian STE20-like protein kinases 1 and 2 (MST1/2), homologs of *Drosophila* Hippo, phosphorylate the effector kinases large tumor suppressors 1 and 2 (LATS1/2), which in turn phosphorylate the transcription factor YES-associate protein (YAP) and sequester it in the cytoplasm^13,14^. The loss of cellular contact reduces MST1/2 and LATS1/2 activity, enabling YAP to enter the nucleus and bind TEA-domain transcription factor (TEAD) to activate pro-proliferative target genes^15,16^. Recent studies have shown that expressing activated YAP and deleting salvador, a scaffold protein needed for MST1/2 activity, are both sufficient to promote adult CM proliferation and myocardial regeneration^17–19^. Despite these findings, the upstream receptor-mediated signals that control the Hippo/YAP signaling are not fully understood.

WNT signaling is well known to be crucial for cardiac development^20^, however a precise role in adult myocardial biology is less clear. The Frizzled (FZD) family of G-protein coupled receptors transduces signaling by WNT ligands^21,22^. These secreted glycoproteins control cell fate, proliferation, and survival via a canonical pathway that prevents a complex containing glycogen synthase kinases α and β (GSK3αβ), axin and adenomatous polyposis coli from phosphorylating the adhesion protein β-catenin^23–25^. Canonical WNT signaling thus stabilizes β-catenin and allows it to enter the nucleus, where it binds T-cell factor (TCF) transcription factors to activate gene expression^26^. In non-cardiac cells, YAP binds axin and is needed for GSK3αβ to phosphorylate β-catenin^27^. YAP is concomitantly degraded with β-catenin and released with β-catenin when GSK3αβ are inhibited^27^. Moreover, β-catenin and YAP bind in the nuclei of tumor cells and embryonic CMs to cooperatively induce the transcription of pro-proliferative targets^28,29^. The canonical Wnt and Hippo signaling pathways are thus integrated into a broader regulatory network.

Studies of WNT signaling in the adult heart have examined β-catenin function in the context of experimental models of MI or hypertrophy, with mixed results reporting that β-catenin protects or exacerbates murine cardiac function depending on the study design^30–32^. Previous studies have also shown that several WNT antagonists of the secreted frizzled-related protein (SFRP) family are cardioprotective against injury using mouse models of MI^33,34^, while downregulation of the WNT/FZD co-receptor LRP6 has been shown to induce adult CM proliferation and regeneration after surgically induced MI in mice^35^. In addition to signaling through the canonical WNT pathway, binding between a subset of WNT ligands and FZD receptors regulates cell polarity, adhesion, and motility through β-catenin-independent pathways^36–38^. This non-canonical WNT signaling also promotes the differentiation of many cell types by attenuating canonical WNT/β-catenin-dependent TCF transcription^39–44^. While numerous studies have examined how canonical WNT signaling affects adult CMs^31,45–50^, the roles for non-canonical WNT signaling in the adult heart remain largely unexplored. Overall, the complex nature of WNT signaling (and its potential crosstalk with other signaling pathways^51^) has prevented our full understanding of how WNT/FZD interactions precisely function in cardiac biology. Comprehensive empirical interrogation of specific ligand and/or receptor function is likely needed before a clear role for WNT signaling in the adult heart can be established.

FZD2, a non-canonical WNT receptor, has been shown to antagonize β-catenin in certain contexts^52–54^. Interestingly, FZD2 was reported to be upregulated in mice after surgically induced MI^33^, suggesting a functional role for this receptor in regulating adult cardiac homeostasis. To investigate this hypothesis, we deleted a conditional allele of *Fzd2* (*Fzd2^floxed^*)^55^ from the CMs of adult mice to identify roles for FZD2 in heart function and repair. The data presented herein reveal that FZD2-dependent WNT signaling prevents CM proliferation and attenuates YAP activity. Furthermore, we find that murine CM-specific FZD2-depletion promotes a cardioprotective and/or reparative response after MI, and thus provide evidence that inhibiting FZD2 may be a novel target pharmacological interventions after MI.

## MATERIALS AND METHODS

### Mouse breeding and genetics

*Fzd2*^floxed^ mice^55^ were mated to *Myh6^CreERT2^* mice^56^ to produce *Fzd2^+/+^; Myh6^CreERT2^* (*Fzd2* WT), *Fzd2^flox/+^; Myh6^CreERT2^* (*Fzd2* HET) and Fzd2^flox/flox^; Myh6*^CreERT2^* (*Fzd2* CKO) mice. *Fzd2^floxed^* mice were bred to *TCF/LEF-GFP* mice^57^ to generate *Fzd2* HET and *Fzd2* CKO mice carrying the transgenic reporter. All alleles were maintained on a mixed CD1 background for at least 5 generations prior to the study. At 8 weeks of age, mice received intraperitoneal injections of 30 mg/kg tamoxifen (TAM) in sesame oil for 3 days to activate CreERT2. When indicated, mice received intraperitoneal injections of 50 mg/kg 5-bromo-2’-deoxyuridine (BrdU) in phosphate buffered saline (PBS) every other day for 10 days (5 injections total). BrdU injections began 12 days before sacrifice to examine CM proliferation under basal conditions and 1 day after coronary artery ligation to assess CM proliferation after MI. Studies were approved by the Institutional Animal Care and Use Committee (IACUC) at the University of Rochester before experiments began and monitored by the University Committee of Animal Resources (UCAR) to ensure compliance with applicable rules and guidelines.

### Animal surgeries

Coronary artery ligations were performed according to a protocol adapted from a prior study^58^. Briefly, mice were anesthetized with 2.0% isoflurane, given subcutaneous injections of 0.3 mg/kg buprenorphine, placed on a heated board and intubated with a PE90 tube connected to ventilator supplying oxygen and isoflurane (0.4 ml tidal volume at 130 breaths/min). Incisions were made on the left side of the chest and a 0.5 cm hole opened between the 3^rd^ and 4^th^ ribs. Hearts were “popped out” by placing pressure on the right side of the rib cage. Left descending coronary arteries were ligated with 6-0 sutures placed 3 mm from their origins. Ligations were confirmed by blanching of the anterior wall. Hearts were then placed back into the thoracic space and incisions closed with purse-string sutures.

### *Ex vivo* ischemia-reperfusion

*Ex vivo* ischemia-reperfusion (I/R) was performed as described^59,60^. Briefly, aortas were cannulated *in situ* prior to hearts being removed and suspended on a Langendorff apparatus. A pressure-transducing balloon was inserted into the LV to track rate pressure product (RPP) over time. Hearts were perfused with Krebs-Henseleit buffer for 25 minutes before being subject to 20 minutes of ischemia by stopping the flow of buffer. Flow was restored and LV contraction observed for 60 minutes of recovery. After reperfusion, hearts were cut into 2 mm slices and stained in tetrazolium chloride (TTC) before being fixed and imaged. Segmentation of tissue from background and infarct from healthy tissue was done with custom MATLAB code.

### Additional materials and methods

A complete materials and methods section with descriptions of adult and neonatal CM isolation, siRNA and plasmid transfections, reporter assays, quantitative real-time RT-PCR (Q-PCR), western blotting and echocardiography can be found in the supplemental methods.

## RESULTS

### Deleting FZD2 from CMs causes cardiomegaly without hypertrophy

*Fzd2* expression was previously shown to be increased in the hearts of mice after MI^33^. To determine if *Fzd2* is induced in CMs or non-myocardial cells such as fibroblasts, which comprise a large portion of the heart’s mass and expand after MI^61^, *Fzd2* mRNA was assessed in CMs and fibroblasts isolated from mice 5 days after permanent ligation of the left descending coronary artery to induce MI. *Fzd2* mRNA levels were higher in CMs from mice subjected to MI than in sham-operated controls (Figure 1A) but unaffected in fibroblasts, suggesting that FZD2 regulates the response of CMs to ischemic injury.

**Figure 1.**
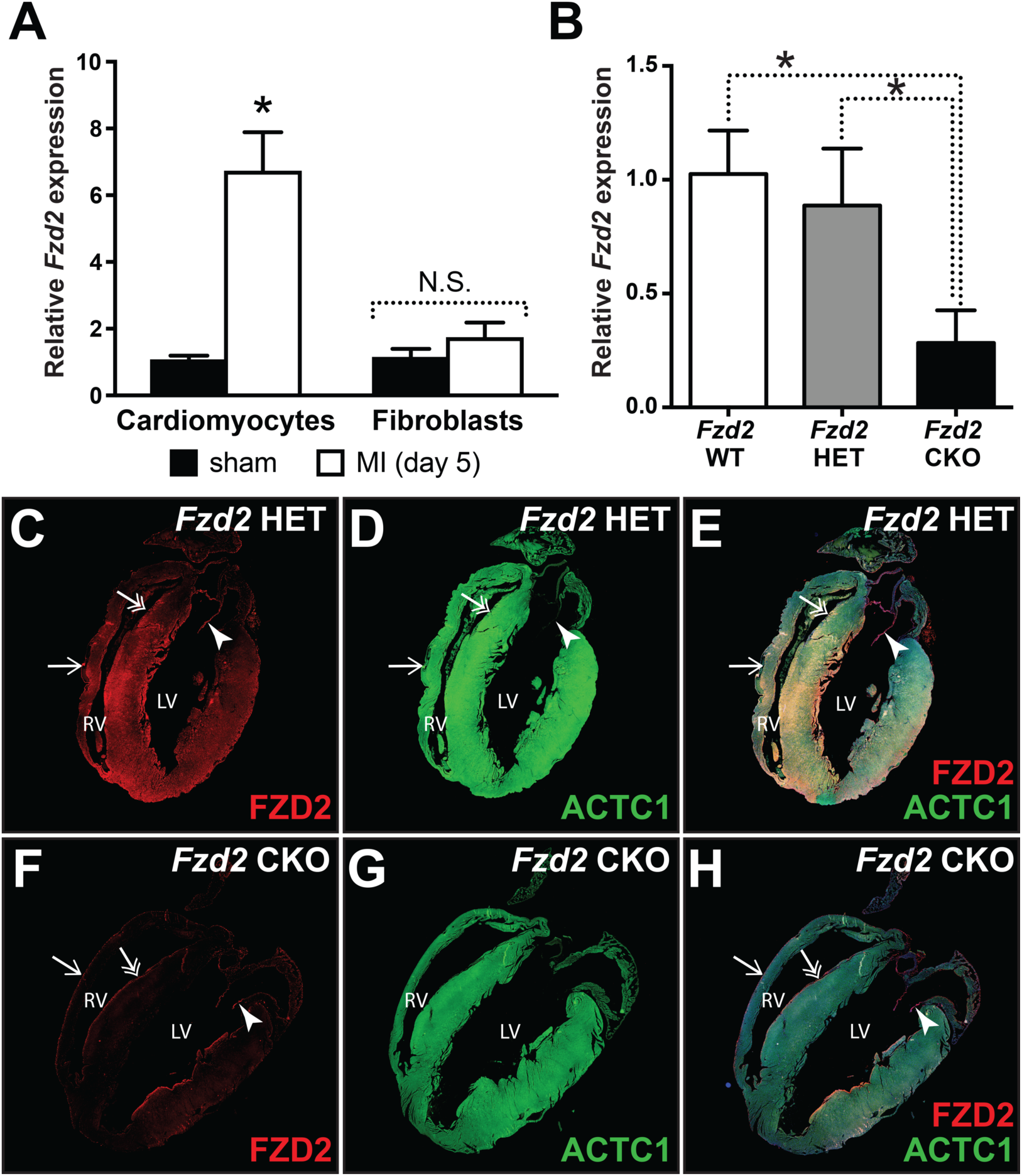
*Fzd2* is expressed in the myocardium of adult mice. **A.** Relative levels of *Fzd2* mRNA in cardiomyocytes (CMs) and fibroblasts from wild type mice 5 days after sham operations or ligation of the left descending coronary artery to induce myocardial infarction (MI). **B.** The relative expression of *Fzd2* in CMs isolated from 4-month-old *Fzd2^+/+^*; *Myh6^CreERT2^* (*Fzd2* WT), *Fzd2^floxed/+^*; *Myh6^CreERT2^* (*Fzd2* HET) and *Fzd2^floxed/floxed^*; *Myh6^CreERT2^* (*Fzd2* CKO) mice injected with tamoxifen (TAM) at 2-months of age. **C-H.** Sections of *Fzd2* HET (C-D) and *Fzd2* CKO (F-H) mice stained for FZD2 (red), ACTC1 (green), and DAPI (blue). Arrows, double headed arrows, and arrowheads point to the epicardium, endocardium, and valves, respectively. Graphs represent the mean ± S.D. for CMs from 4 mice of each genotype. Asterisks indicate p ≤ 0.05 by Student’s *t*-test (A) or one-way ANOVA with Tukey’s multiple comparisons test (B).

To further define the roles FZD2 plays in the adult myocardium, mice carrying *Fzd2^floxed^* were bred to *Myh6^CreERT2^* mice, which express tamoxifen (TAM) inducible Cre in CMs^56^. The resulting *Fzd2^+/+^; Myh6^CreERT2^* (*Fzd2* WT), *Fzd2^floxed/+^; Myh6^CreERT2^* (*Fzd2* HET) and *Fzd2^floxed/floxed^; Myh6^CreERT2^* (*Fzd2* CKO) progeny were injected with TAM at 2 months of age and sacrificed at 4 months of age. Hearts were harvested and perfused with collagenase to isolate CMs and produce CM-enriched RNA from each mouse. *Fzd2* mRNA was significantly reduced in *Fzd2* CKO CMs relative to *Fzd2* WT and *Fzd2* HET CMs (Figure 1B). *Fzd2* mRNA levels were not significantly different between *Fzd2* WT and *Fzd2* HET CMs. Immunological staining for FZD2 and the CM-specific isoform of actin (ACTC1) revealed FZD2 expression in CMs of the ventricles and interventricular septa of *Fzd2* HET mice, but not in those of *Fzd2* CKO mice (Figure 1C-H). FZD2 staining was similar in the epicardium (arrows), endocardium (double headed arrows), and valves (arrowheads) of *Fzd2* HET and *Fzd2* CKO mice, consistent with the loss of FZD2 being CM-specific. The *Fzd2^floxed^* allele may be incorrectly targeted resulting in a more complex rearrangement leading to a hypomorph after TAM treatment^62^. Nevertheless, we find strongly reduced *Fzd2* expression in our current study after TAM injections in *Fzd2* CKO hearts relative to controls, and previous work using this allele similarly reported a strong reduction of *Fzd2* expression using different Cre-lines and organ contexts^55,63^. Thus, if a hypomorph for FZD2 protein is generated in CMs in *Fzd2* CKO mice after TAM treatment it is likely non-functional.

The hearts of *Fzd2* HET mice were similar in size to *Fzd2* WT mice after TAM injection (data not shown), and as the *Fzd2* mRNA expression level was also unchanged between *Fzd2* WT and *Fzd2* HET hearts after TAM injection (Figure 1B), we concluded that loss of one copy of *Fzd2* is insufficient to cause a phenotype and thus *Fzd2* HET mice were used interchangeably as controls with *Fzd2* WT (where indicated) in subsequent experiments. We found that *Fzd2* CKO mice had grossly larger hearts and higher heart weight to tibia length ratios than *Fzd2* HET mice after TAM injection (Figure 2A-C). However, *Fzd2* CKO CMs had no significant difference in the cross-sectional areas relative to controls in sectioned hearts stained with wheat germ agglutinin (WGA), which labels cell membranes (Figure 2D-F). The cardiomegaly in *Fzd2* CKO mice was thus not caused by concentric hypertrophy.

**Figure 2.**
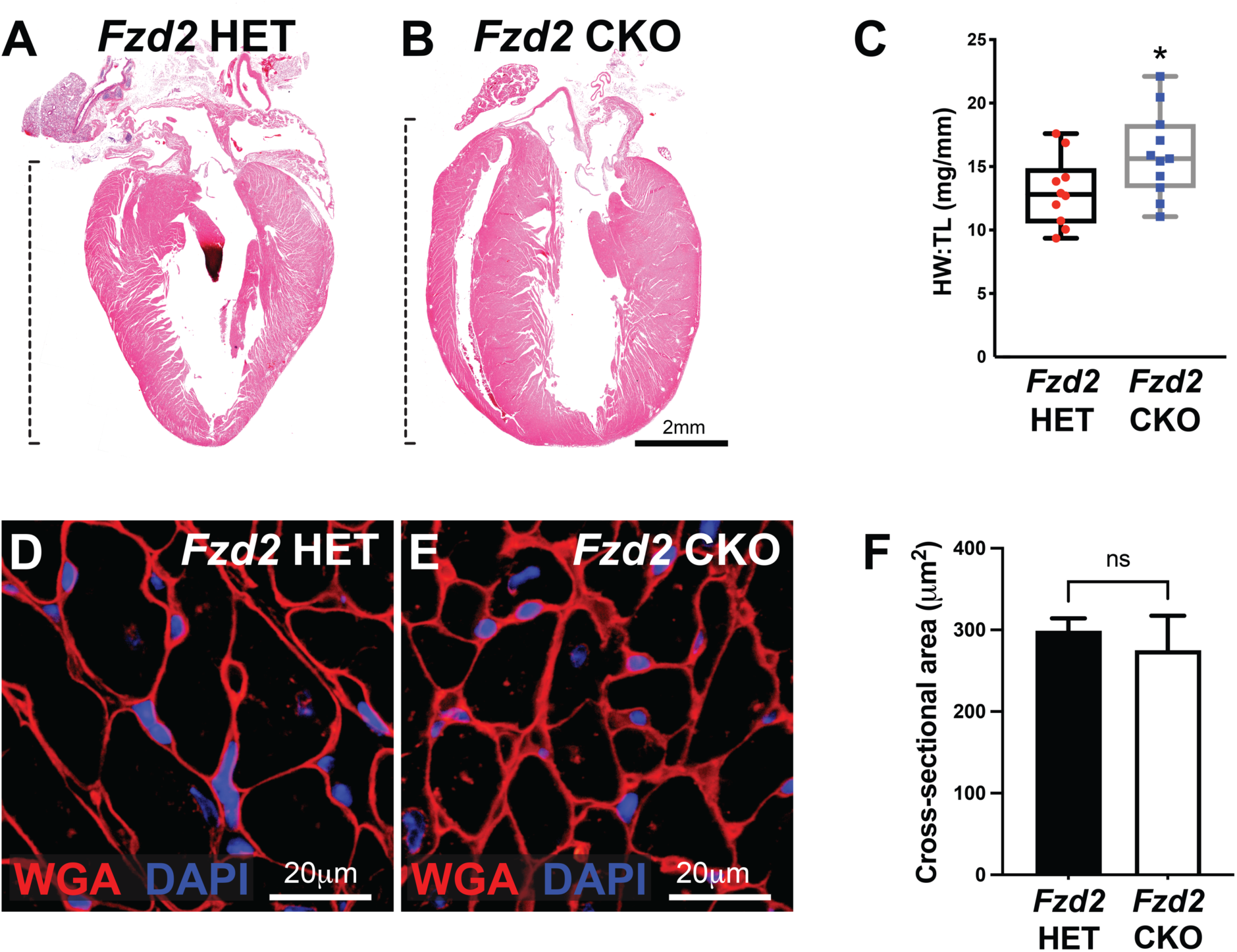
CM-specific FZD2-deletion causes cardiomegaly but not CM hypertrophy. **A, B.** H&E stained sections of hearts from 4-month old *Fzd2* HET (A) and *Fzd2* CKO (B) mice injected with TAM at 2-months of age. **C.** Graph shows mean heart weight to tibia length (HW:TL) ratios for *Fzd2* HET (n=10) and *Fzd2* CKO (n=11) mice. Box plot shows minimum, first quartile, median, third quartile, and maximum values. **D, E.** Sectioned hearts of *Fzd2* HET (D) and *Fzd2* CKO (E) mice stained with wheat germ agglutinin (WGA, red) and DAPI (blue). **F.** Graph shows the mean ± S.E.M. cross-sectional area of CMs in *Fzd2* HET (n=3) and *Fzd2* CKO (n=3) mice. Asterisks indicate p ≤ 0.05 by Student’s *t*-tests.

### Deleting FZD2 from adult CMs induces cell cycle reentry

To determine if *Fzd2* CKO hearts were larger due to increased CM proliferation, *Fzd2* CKO and *Fzd2* HET mice were injected with TAM, recovered for 1 month, and then given 5 injections of BrdU at 48-hour intervals. Mice were sacrificed 48 hours after the last injection to collect hearts for sectioning. Sections were stained for WGA, DAPI, and antibodies for BrdU and Desmin, which localizes to the Z-bands of CMs. Most BrdU+ nuclei were small and located outside CM membrane boundaries in *Fzd2* HET mice and likely represent non-CM cell types (arrowheads, Figure 3A, C). In contrast, BrdU+ nuclei were identified clearly within a small proportion of CMs in *Fzd2* CKO hearts at a significantly higher frequency than in *Fzd2* HET controls (Figure 3B, D, E). Isolated CMs from *Fzd2* WT and *Fzd2* CKO mice were also stained for DAPI, cardiac Troponin T (TNNT2), and the proliferation marker Ki67 (Figure 3F-I). We were unable to find Ki67+ cells among *Fzd2* WT CMs. Nevertheless, a significant 0.58% of CMs from *Fzd2* CKO mice had Ki67+ nuclei (Figure 3J).

**Figure 3.**
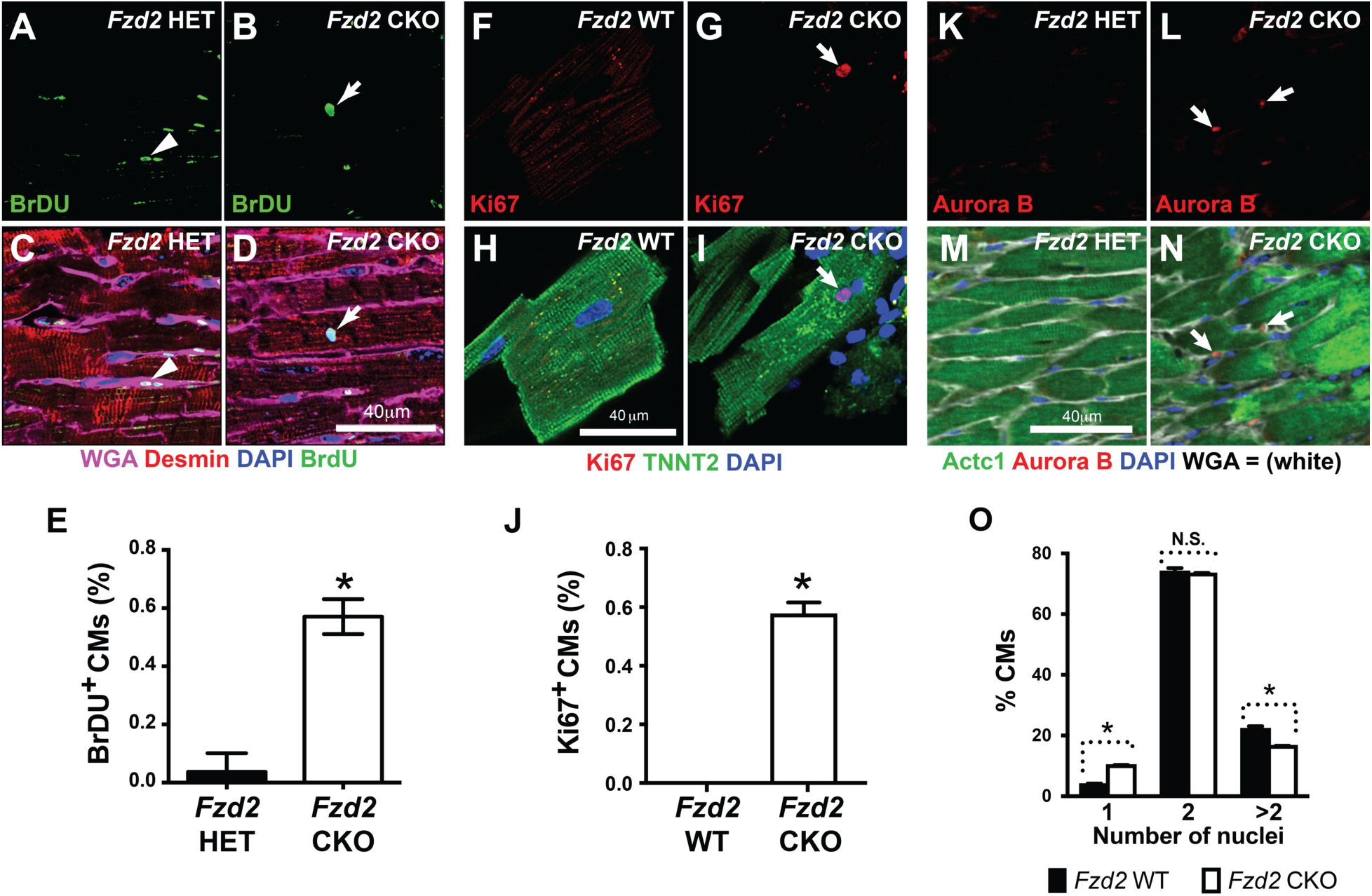
FZD2 inhibits CM proliferation in adult mice. **A-D.** Sections of *Fzd2* HET (A, C) and *Fzd2* CKO (B, D) mice stained with WGA (violet), DAPI (blue) and antibodies for BrdU (green) and Desmin (red). Arrows in (B) and (D) show the BrdU+ nucleus of a CM. Arrowheads in (A) and (C) show BrdU+ non-CM cells between CMs. **E.** Mean ± S.E.M. percentages of Desmin+ CMs with BrdU+ nuclei in *Fzd2* HET (n=3) and *Fzd2* CKO (n=3) mice. **F-I.** CMs isolated from adult *Fzd2* WT and *Fzd2* CKO mice stained for DAPI (blue), Ki67 (red), and TNNT2 (green). Arrows in (G) and (I) point to a Ki67+ CM nucleus. **J.** Mean ± S.E.M. percentage of isolated CMs with Ki67+ nuclei in *Fzd2* WT (n=3) and *Fzd2* CKO (n=3) mice. **K-N.** Sections of *Fzd2* HET and *Fzd2* CKO hearts stained with DAPI (blue), WGA (white) and antibodies for aurora b (red), and ACTC1 (green). Arrows in (L) and (N) show aurora b staining between CMs. **O.** Mean ± S.D. percentage of CMs from *Fzd2* WT (n=3) and *Fzd2* CKO (n=3) mice with one, two, or more than two nuclei. Asterisks indicate p ≤ 0.05 on student’s t-test (E, J) or two-way ANOVA with Tukey’s multiple comparisons test (O).

Since most CMs in adult mice are binucleated, the incorporation of BrdU into the nuclei of *Fzd2* CKO CMs may reflect DNA replication without cytokinesis. Sectioned hearts were thus stained for WGA, DAPI, ACTC1, and aurora b, a kinase that localizes to the cleavage furrows of cells undergoing cytokinesis^64–66^. Aurora b was not found in the ACTC1 labeled CMs of *Fzd2* HET mice (Figure 3K, M). In contrast, aurora b staining was occasionally observed at punctate regions of membrane between *Fzd2* CKO CMs (arrows, Figure 3L, N), suggesting the DNA synthesis in these cells leads to cytokinesis and the production of new CMs. Moreover, if the CMs of *Fzd2* CKO mice replicated their DNA without dividing an increased number of CMs would have two or more nuclei relative to controls. However, *Fzd2* CKO CMs were more likely to be mono-nucleated than those of *Fzd2* WT controls (Figure 3O), further suggesting that some *Fzd2* CKO CMs undergo cytokinesis.

### FZD2-deletion increases β-catenin protein levels but not TCF-dependent transcription

Consistent with reports of FZD2 inhibiting canonical WNT signaling^52^, β-catenin protein was enriched in whole heart lysates from *Fzd2* CKO mice relative to those of *Fzd2* HET mice (Figure 4A). Sectioned hearts from *Fzd2* CKO and *Fzd2* HET mice were stained for WGA, DAPI, β-catenin, and the desmosome component desmoplakin. In *Fzd2* HET mice β-catenin and desmoplakin co-localized at intercalated discs, specialized junctions between adjacent CMs (arrows, Figure 4B-D). In contrast, β-catenin staining was stronger at the intercalated discs (arrows, Figure 4E-G) and present along the lateral membranes of *Fzd2* CKO CMs (arrowheads, Figure 4E-G). Yet, while β-catenin levels were higher in *Fzd2* CKO CMs than controls, β-catenin was not enriched in the nuclei of *Fzd2* CKO CMs and may not activate TCF-dependent transcription in these cells.

**Figure 4.**
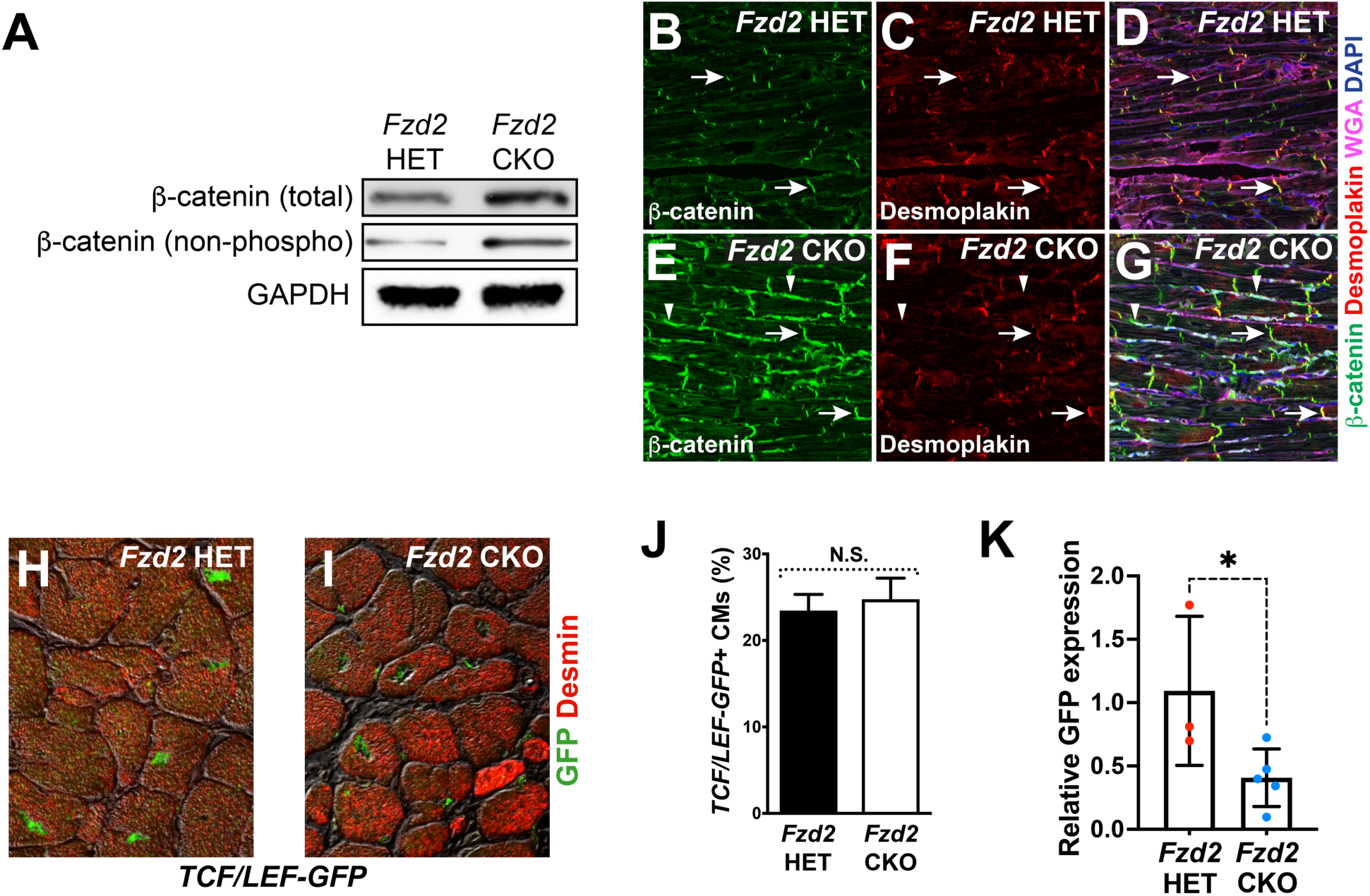
FZD2 loss-of-function increases the levels of β-catenin protein but not TCF-dependent transcription. **A.** Western blots for total β-catenin, non-phospho-β-catenin and GAPDH (loading control) in whole hearts of *Fzd2* HET and *Fzd2* CKO mice. **B-G.** Sections of *Fzd2* HET (B, C, D) and *Fzd2* CKO (E, F, G) hearts stained for β-catenin (green), desmoplakin (red), WGA (violet), and DAPI (blue). Arrows indicate staining at intercalated discs. Arrowheads (E, F, G) indicate β-catenin staining at the lateral edges of *Fzd2* CKO CMs. **H, I.** Sections of *Fzd2* HET (H) and *Fzd2* CKO (I) hearts carrying the *TCF/LEF-GFP* reporter stained for GFP (green) and Desmin (red). Staining is layered over DIC images to enhance contrast. **J.** Graph shows the mean ± S.E.M. percentage of Desmin+ CMs expressing *TCF/LEF-GFP* in *Fzd2* HET (n=2) and *Fzd2* CKO (n=3) mice. **K.** Relative expression ± S.D. of *GFP* in CMs from *Fzd2* HET (n=3) and *Fzd2* CKO (n=5) hearts. N.S. indicates non-significant, asterisk indicates p ≤ 0.05 by one-tailed Student’s *t*-test.

To determine if the excess β-catenin in FZD2-deficient CMs induces TCF-dependent transcription, *Fzd2* CKO and *Fzd2* HET mice carrying *TCF/LEF-GFP* were generated. This reporter uses tandem repeats of the consensus TCF binding site to express nuclear GFP in cells receiving canonical WNT signaling. Surprisingly, co-staining sectioned hearts for GFP and desmin revealed *TCF/LEF-GFP* activity in equal numbers of CMs in *Fzd2* CKO and *Fzd2* HET mice (4H-J). However, when the expression of GFP was examined by RT-qPCR using RNA isolated from the hearts of Fzd2 CKO and Fzd2 HET mice we found GFP expression to be significantly decreased in Fzd2 CKO hearts relative to Fzd2 HET controls (Figure 4K). Together, these results suggest that Fzd2 loss-of-function increases β-catenin protein levels in CMs at the expense of TCF/LEF transcriptional activity.

### FZD2-deletion increases YAP protein and target gene expression *in vivo*

The CM proliferation in adult *Fzd2* CKO mice is strikingly reminiscent of the effects of disrupting Hippo signaling^17–19,29^. FZD2 may thus prevent adult CM proliferation by restraining YAP in the adult myocardium. Consistent with this idea, staining for YAP was more intense in the nuclei of CMs isolated from adult *Fzd2* CKO mice than controls (Figure 5A, B). Western blotting confirmed that YAP protein was enriched in the hearts of *Fzd2* CKO mice relative to controls (Figure 5C, D). The phosphorylation of YAP at serine residues 127 and 397, which cause nuclear exclusion^67^ and degradation^68^, respectively, were also reduced in *Fzd2* CKO hearts (Figure 5C, E, F). YAP target gene expression was also increased in the CMs of *Fzd2* CKO mice relative to controls, including the proto-oncogene *v-myc avian myelocytomatosis viral oncogene 1* (*Mycl*, Figure 5G) and the apoptosis regulator *B cell leukemia/lymphoma 2* (*Bcl2l1,* Figure 5H). Increased YAP activity may thus contribute to the cardiomegaly and increased CM cell cycle activity in *Fzd2* CKO mice.

**Figure 5.**
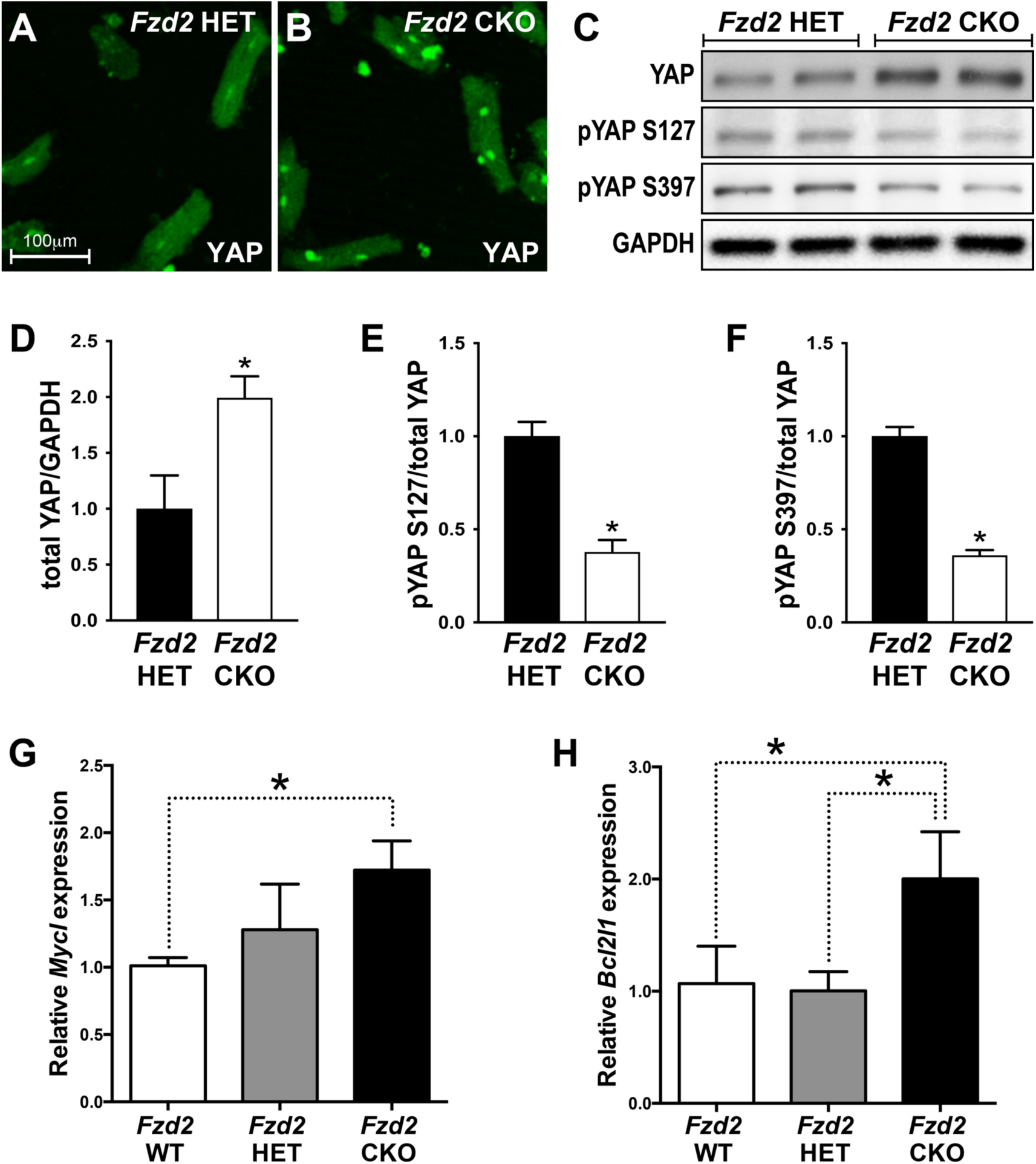
FZD2 loss-of-function increases the levels of YAP protein and targets *in vivo*. **A,B.** CMs from *Fzd2* HET (A) and *Fzd2* CKO (B) mice stained for YAP (green). **C.** Western blots for total YAP, and YAP phosphorylated at serine 127 (pYAP S127) and 397 (pYAP S397), and GAPDH in whole hearts of *Fzd2* HET and *Fzd2* CKO mice. **D.** Relative levels of YAP protein normalized to GAPDH in the hearts of *Fzd2* HET and *Fzd2* CKO mice. **E, F.** Relative levels of pYAP S127 (E) and pYAP S397 (F) normalized to total YAP in *Fzd2* HET and *Fzd2* CKO hearts. Graphs in D-F show mean intensity ± S.E.M. of bands from lysates of *Fzd2* HET (n=3) and *Fzd2* CKO (n=3) mouse hearts. **G, F.** Relative expression ± S.D. of *Mycl* (G), and *Bcl2l1* (H) in CMs isolated from *Fzd2* WT (n=4) and *Fzd2* CKO (n=4) mouse hearts. Asterisks indicate p ≤ 0.05 by either Student’s t-test (D-F) or one-way ANOVA with Tukey’s multiple comparisons test (G, H).

### FZD2 knockdown increases YAP protein levels in cultured neonatal CMs

Neonatal ventricular CMs (NVCMs) were used to further examine the effects of FZD2 on YAP *in vitro*. Transfecting NVCMs with siRNA targeting *Fzd2* caused ∼95% knockdown (KD) of *Fzd2* expression relative to controls 72 hours after transfection (Figure 6A). Consistent with the elevated levels of YAP observed in *Fzd2* CKO CMs, YAP protein was increased in NVCMs transfected with FZD2 siRNA relative to controls (Figure 6B). YAP staining was also stronger in NVCMs transfected with FZD2 siRNA (Figure 6F-H) than in control siRNA treated NVCMs (Figure 6C-E). Quantification of the YAP fluorescence signal intensity from stained NVCMs confirmed an increase in whole-cell and nuclear-specific YAP levels in FZD2 KD NVCMs relative to controls (Figure 6O, P). Together, these data suggest that FZD2 deficiency increases YAP activity by stabilizing the YAP protein.

**Figure 6.**
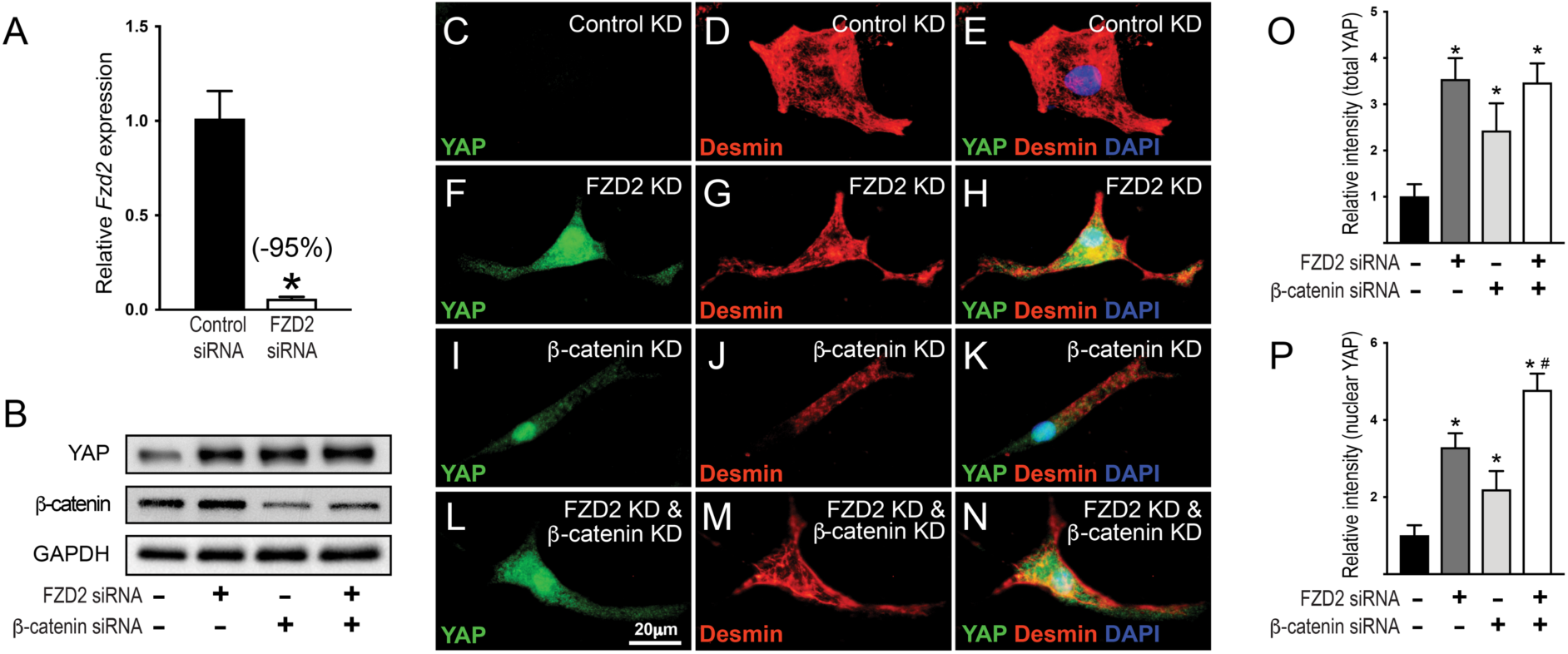
FZD2-deficiency increases YAP protein expression via a β-catenin-independent mechanism. **A.** Relative expression of *Fzd2* in neonatal ventricular CMs (NVCMs) transfected with scrambled (control) or FZD2 siRNA. **B.** Western blots for YAP, β-catenin and GAPDH in NVCMs transfected with FZD2 and β-catenin siRNA. **C-N.** NVCMs transfected with control siRNA (C-E), FZD2 siRNA (F-H), β-catenin siRNA (I-K) or both FZD2 and β-catenin siRNA (L-N) stained with DAPI (blue) and antibodies for YAP (green) and Desmin (red). **O, P.** Relative intensities of total (O) and nuclear YAP (P) staining in NVCMs transfected FZD2 and β-catenin siRNA as in **C-N**. Graphs represent the mean ± S.D, asterisks indicate p ≤ 0.05 relative to scrambled siRNA control, # indicates p ≤ 0.05 relative to all other sample comparisons, by one-way ANOVA with Tukey’s multiple comparisons test.

Prior studies showed that β-catenin binds YAP and enhances its induction of pro-proliferative targets in tumor cells and embryonic CMs^29^, implying that the high levels of β-catenin in FZD2-deficient CMs may activate YAP. NVCMs were thus transfected with FZD2 and β-catenin siRNA, alone or in combination, to determine if β-catenin is required for the increased levels of YAP in FZD2 KD NVCMs. Western blotting confirmed that β-catenin was reduced in NVCMs transfected with β-catenin siRNA relative to controls (Figure 6B). Surprisingly, β-catenin KD increased YAP protein in NVCMs relative to controls when assessed by western blotting and immunological staining to levels comparable to those in FZD2 KD NVCMs (Figure 6B, I-K, O, P). Co-transfecting NVCMs with FZD2 and β-catenin siRNA did not increase total YAP levels above those in NVCMs treated with either siRNA alone (Figure 6B, L-N, O). However, NVCMs treated with both FZD2 and β-catenin siRNA had higher levels of nuclear YAP than those treated with either siRNA alone (Figure 6O).

### FZD2 knockdown increases YAP reporter activity in cultured neonatal CMs

To examine the effects of FZD2 on YAP activity *in vitro*, NVCMs were transfected with FZD2 siRNA and a two-component reporter consisting of TEAD1 fused to the GAL4 DNA-binding domain and a UAS-driven luciferase^15^. While TEAD1-reporter activity trended higher in NVCMs treated with FZD2 siRNA relative to controls, the signal from endogenous YAP was too weak and noisy to observe significant changes. NVCMs were thus co-transfected with a YAP1 expression vector to increase the signal in our assays. As expected, YAP1 increased TEAD1-reporter activity ∼12x in NVCMs transfected with control siRNA (Figure 7A). In contrast, YAP1 increased reporter activity ∼17x in NVCMs transfected with FZD2 siRNA, a level ∼44% greater than observed control siRNA treated NVCMs (Figure 7A). A similar result was observed when transfecting with 8x-TEAD-Luc, a YAP reporter that uses 8 copies of the consensus TEAD binding site to drive luciferase expression^69^, where YAP1 expression increased reporter activity to a greater degree in NVCMs transfected with FZD2 siRNA relative to control siRNA (Figure 7B). In contrast, YAP1-dependent TEAD1-reporter activity was reduced ∼49% in NVCMs transfected with a FZD2 expression plasmid compared to empty vector controls (Figure 7C). FZD2 is thus necessary and sufficient to inhibit YAP-dependent transcription in CMs.

**Figure 7.**
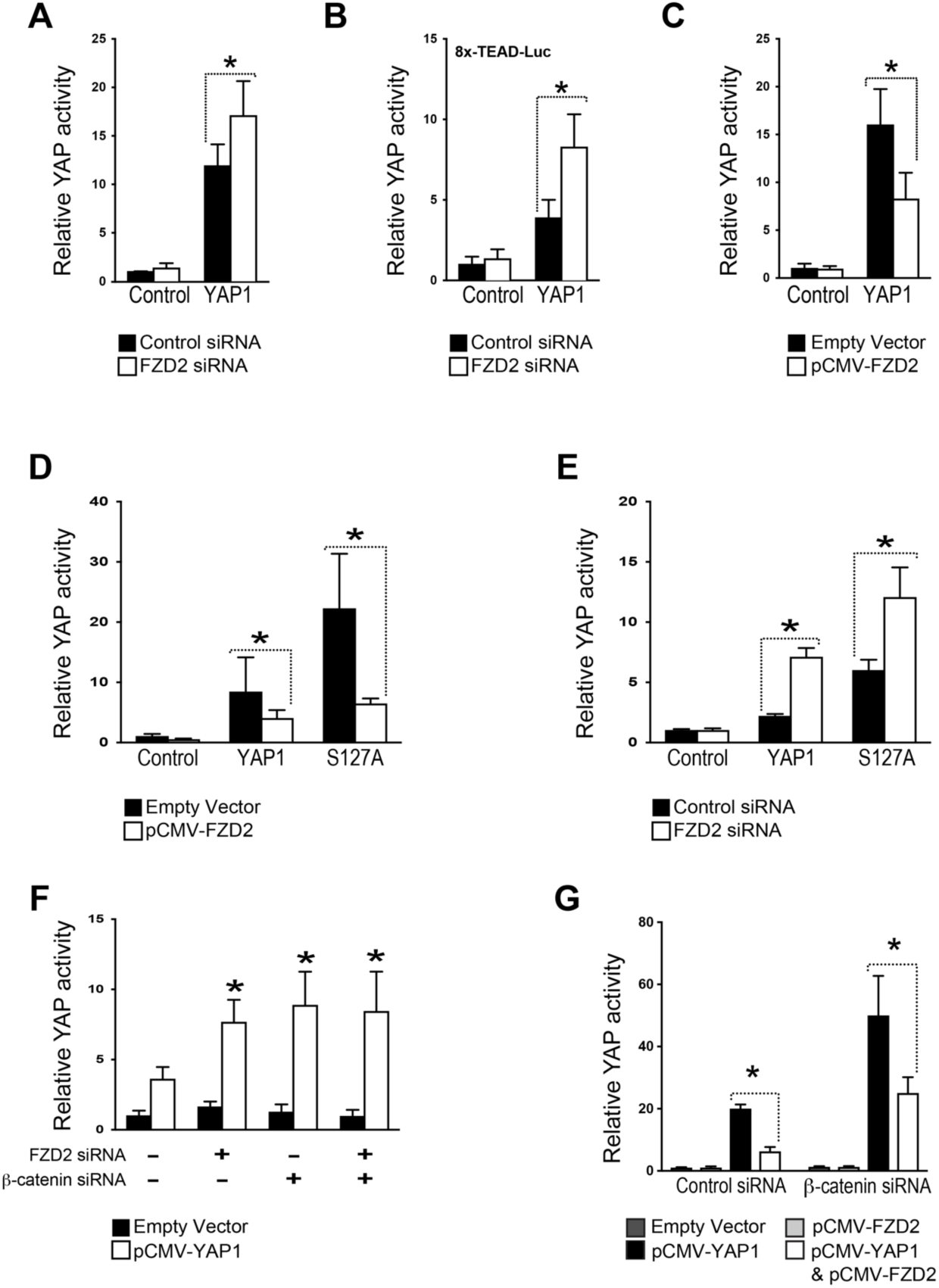
FZD2-deficiency increases YAP reporter activity via a β-catenin-independent mechanism. **A.** Relative activity of TEAD-Gal4-UAS-Luc, a YAP reporter consisting of TEAD1 fused to the GAL4 DNA-binding domain and a UAS-driven firefly luciferase, in NVCMs transfected with empty vector or YAP1 expression plasmid and either control or FZD2 siRNA. **B.** Relative activity of the 8x-TEAD-Luc reporter, which uses 8 copies of the TEAD consensus binding site to express firefly luciferase, in NVCMs transfected with empty vector or YAP1-expression plasmid and control or FZD2 siRNA. **C.** TEAD-Gal4-UAS-Luc activity in NVCMs transfected with empty vector or YAP expression plasmid with or without FZD2 expression vector. **D.** TEAD-Gal4-UAS-Luc activity in NVCMs transfected with empty vector, YAP-expression vector, or expression plasmid for a YAP mutant with serine 127 mutated to alanine (YAP S127A) either alone or in combination with FZD2 expression vector. **E.** TEAD-Gal4-UAS-Luc activity in NVCMs transfected with empty vector, YAP expression plasmid, or YAP S127A expression vector and either control or FZD2 siRNA. **F.** TEAD-Gal4-UAS-Luc activity in NVCMs transfected with empty vector or YAP1 expression plasmid and FZD2 or β-catenin siRNA. **G.** TEAD-Gal4-UAS-Luc activity in NVCMs transfected with empty vector, YAP1 expression vector, FZD2 expression vector, or both FZD2 and YAP1 vectors as well as control or β-catenin siRNA. Graphs represent the mean ± S.D, asterisks indicate p ≤ 0.05 by two-way ANOVA with Tukey’s multiple comparisons test.

To determine if the effects of FZD2 require phosphorylation by LATS1/2, NVCMs were co-transfected with the TEAD1-reporter and an expression plasmid for YAP-S127A, a constitutively active YAP mutant with the LATS1/2 target site at serine 127 mutated to alanine (S127A). YAP-S127A increased TEAD1-reporter activity ∼2.7x more than wild type YAP1 in NVCMs (Figure 7D) consistent with data indicating that S127 phosphorylation inhibits YAP by excluding it from the nucleus^67^. However, co-transfecting NVCMs with FZD2 and YAP1-S127A expression plasmids reduced TEAD1-reporter activity ∼71% relative to NVCMs expressing YAP1-S127A alone (Figure 7D). Moreover, TEAD1 reporter activity was ∼2x higher when NVCMs were transfected with YAP-S127A plasmid and FZD2 siRNA relative to those transfected with YAP-S127A plasmid and control siRNA (Figure 7E). FZD2-mediated YAP inhibition is thus independent of serine 127 phosphorylation.

To determine if β-catenin is necessary for FZD2 KD to increase YAP activity, NVCMs were transfected with the TEAD1-reporter and siRNA targeting β-catenin and FZD2, alone or in combination. Consistent with the effects of β-catenin KD on YAP protein, co-transfecting NVCMs with β-catenin siRNA increased the levels of YAP-dependent reporter activity above those in NVCMs transfected with control siRNA (Figure 7F). Combined KD of FZD2 and β-catenin did not activate YAP more than the KD of either factor alone (Figure 7F). However, FZD2 overexpression inhibited YAP in β-catenin KD NVCMs (Figure 7G), suggesting that FZD2 and β-catenin inhibit YAP through partially independent mechanisms.

### FZD2-deletion improves cardiac function after myocardial infarction

Mutations that cause adult CM proliferation also promote myocardial regeneration after MI^17–19,29^. To determine if deleting FZD2 from adult CMs improves cardiac function after ischemic injury, *Fzd2* HET and *Fzd2* CKO mice were subject to experimental MI 3 months after TAM injection. Echocardiography was performed 1 day prior to MI to establish baseline readings and then repeated 3, 7, 14 and 28 days after MI to follow cardiac function during their recovery. Baseline measurements for LV ejection fraction (EF) and fractional shortening (FS) were equivalent in *Fzd2* HET and *Fzd2* CKO mice (Figure 8A, B, Supplemental Table 3), indicating that FZD2 is not essential for basal cardiac function. However, while EF and FS percentages were significantly reduced in *Fzd2* HET mice at all time points after MI, cardiac function was not significantly reduced in *Fzd2* CKO mice at any time-point after MI (Figure 8A, B). *Fzd2* HET mice also showed other signs of heart failure after MI, including increased LV internal diameter and decreased LV anterior wall thickness, that were not detected in *Fzd2* CKO mice (Supplemental Table 3). To determine if deleting FZD2 from CMs reduced fibrotic scarring, hearts were harvested at the end of the 28-day recovery post-MI, sectioned, and stained with Masson’s trichrome (Figure 8C). The percentage of total area occupied by scar in sections taken every 50 µm from the apex of the heart to the ligature was significantly reduced in the *Fzd2* CKO hearts compared to the controls (Figure 8D).

**Figure 8.**
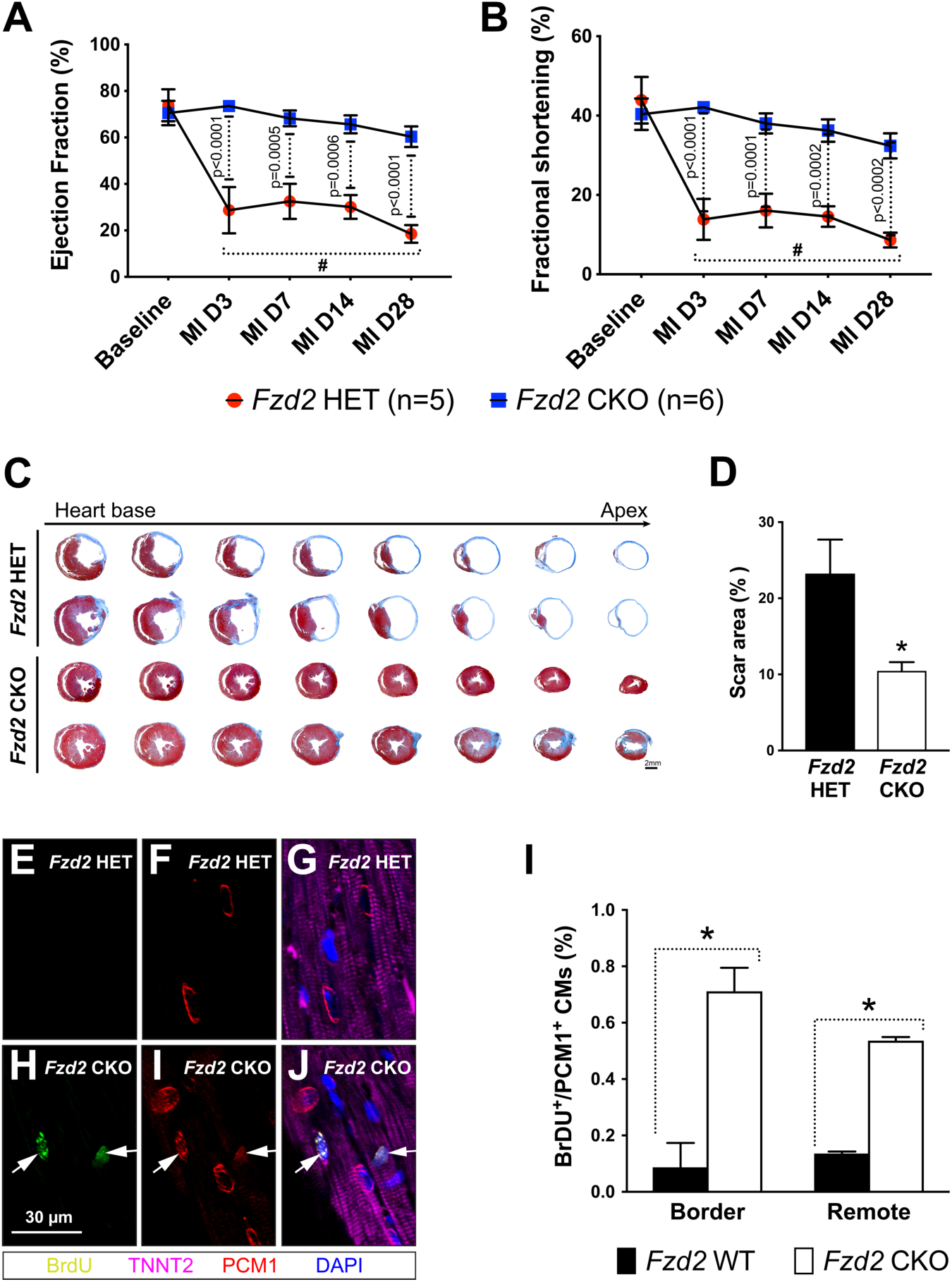
FZD2-deletion improves cardiac function after MI. **A, B.** Graph shows mean ± S.E.M. ejection fraction (**A**) and fractional shortening (**B**) of 5 male *Fzd2* HET and 6 male *Fzd2* CKO mice injected with TAM at 2 months of age and subjected to MI at 5 months of age. Measurements were recorded one day prior to MI (baseline) as well as 3, 7, 14 and 28 days after MI. All echocardiography data are shown in Supplemental Table 3. **C.** Sectioned hearts of *Fzd2* HET and *Fzd2* CKO mice collected 28 days post-MI and stained with Masson’s trichrome. **D.** Graph shows the mean ± S.E.M. percentage of total area covered by scar tissue in the sectioned hearts of *Fzd2* HET and *Fzd2* CKO mice shown in (C). **E-H.** Sectioned hearts of *Fzd2* HET (E, G) and *Fzd2* CKO (F, H) mice collected 28 days after MI and stained with DAPI (blue) and antibodies for BrdU (green), PCM1 (red), and TNNT2 (violet). Arrows point to BrdU+/PCM1+ nuclei of TNNT2+ CMs of *Fzd2* CKO mice. **I.** Graph shows mean ± S.D. percentage of PCM1+/TNNT2+ CMs with BrdU+ nuclei in the hearts of *Fzd2* HET (n=3) and *Fzd2* CKO (n=3) mice 28 days post-MI. Asterisks indicate p ≤ 0.05 by two-way ANOVA with Tukey’s multiple comparisons test (A, B) or Student’s *t*-test (D, I). # indicates p ≤ 0.0002 for *Fzd2* HET data post-MI relative to *Fzd2* HET and *Fzd2* CKO baseline data by two-way ANOVA with Tukey’s multiple comparisons test (A, B).

Remarkably, cardiac function of *Fzd2* CKO mice was not reduced 3 days after MI, the earliest time examined. To determine if this strong cardioprotective effect was due to changes in the vasculature that prevent ischemia, *Fzd2* HET and *Fzd2* CKO mice were subject to MI and recovered for 24 hours before having their hearts perfused with fluorescent microspheres. Perfused hearts were cut into 1 mm slices and stained with TTC, which turns red in living cells after being reduced by NADH and other electron donors in the cytoplasm but remains white in necrotic cells due to the depletion of these co-enzymes^70^, and imaged (Supplemental Figure 1A). Surprisingly, the percentage of myocardium cut off from coronary blood flow or area at risk (AAR) was unaffected by FZD2 loss-of-function (Supplemental Figure 1B). The percentages of myocardium and AAR that turned necrotic, as judged by the lack of TTC staining, were also equivalent in *Fzd2* CKO and *Fzd2* HET hearts (Supplemental Figure 1C, D). The improved outcomes of *Fzd2* CKO mice after MI are thus not due to changes in coronary blood flow.

To further examine the effects of FZD2-deletion on CM survival, the hearts of *Fzd2* HET and *Fzd2* CKO mice were assessed in an *ex vivo* model of ischemia/reperfusion (I/R). Hearts were cannulated, suspended on a Langendorff apparatus and perfused with buffer containing oxygen and nutrients. After having pressure-transducing balloons inserted into their left ventricles, hearts were equilibrated for 25 minutes before being subjected to 20 minutes of ischemia by cutting off the flow of buffer. The flow of buffer was then restored, and the force of LV contraction monitored for 1 hour after reperfusion. The improvement of rate pressure product (RPP) over time was equivalent in *Fzd2* HET and *Fzd2* CKO hearts (Supplemental Figure 1E). TTC staining further showed that the extent of necrosis was similar in the hearts of *Fzd2* CKO and *Fzd2* HET mice (Supplemental Figure F, G). Based on these data, it seems unlikely that the improved cardiac function in *Fzd2* CKO mice after MI results from increased acute post-injury CM survival.

We next determined whether the increased proliferative capacity of FZD2-deficient CMs might contribute to the maintenance of post-MI cardiac function in *Fzd2* CKO mice. *Fzd2* CKO and control mice were injected with BrdU starting 1 day after MI and harvested 28 days later. Sectioned hearts were stained for DAPI, BrdU, TNNT2, and pericentriole material 1 (PCM1), a centriolar satellite protein that localizes to a ring around the nuclei of CMs but not in endothelial cells or fibroblasts (Figure 8E-H). A significantly increased percentage of TNNT2+ and PCM1+ CMs with BrdU+ nuclei were observed in both the border and remote zones of *Fzd2* CKO hearts relative to *Fzd2* HET hearts (Figure 8I). CM proliferation may thus contribute to the improved outcomes of *Fzd2* CKO mice after MI relative to *Fzd2* HET controls.

## DISCUSSION

*Fzd2* CKO mice had larger hearts than *Fzd2* HET mice with equivalent CM cross-sectional areas, therefore the cardiomegaly in *Fzd2* CKO mice is not likely due to concentric hypertrophy. Instead, the CMs of *Fzd2* CKO mice had higher rates of BrdU incorporation and Ki67 expression than controls, suggesting that FZD2 prevents adult CMs from reentering the cell cycle. Aurora b was also present at membranes between CMs in *Fzd2* CKO mice, consistent with these cells undergoing cytokinesis. Moreover, *Fzd2* CKO mice had more mono-nucleated CMs than controls, implying that the cell cycle activity after *Fzd2*-deletion produces new CMs. Based on these data, we conclude that FZD2 normally prevents adult CM proliferation and suggest that the cardiomegaly in *Fzd2* CKO mice is due to CM hyperplasia.

The hearts of *Fzd2* CKO mice contained higher levels of β-catenin than controls. Yet, while β-catenin was enriched in the lateral membranes and intercalated discs of FZD2 deficient CMs, high levels of β-catenin were not found in the nuclei of these cells. The canonical WNT reporter *TCF/LEF-GFP* was also found to be active in equivalent numbers of CMs in the hearts of *Fzd2* CKO and control mice, although the overall GFP expression was reduced in *Fzd2* CKO hearts relative to *Fzd2* HET controls. *Fzd2* loss-of-function may thus increase β-catenin protein throughout CMs at the expense of TCF-mediated transcription. These data agree with those of a prior study, which showed that a constitutively active β-catenin that lacked the amino-terminal GSK3αβ phosphorylation sites failed to enter the nucleus or activate TCF in adult CMs^47^. The effects of FZD2 deletion on CM proliferation are thus unlikely to result from increased TCF-dependent transcription.

The CM proliferation in *Fzd2* CKO mice resembles the effects of activating the transcription factor YAP, which is inhibited by the Hippo pathway, in the adult heart. We thus reasoned that FZD2 loss-of-function could promote adult CM proliferation by increasing YAP activity. Consistent with this hypothesis, YAP protein was enriched in the nuclei of *Fzd2* CKO CMs and *Fzd2* KD NVCMs relative to controls. Moreover, the YAP target genes *Mycl*, and *Bcl2l1* were expressed at higher levels in *Fzd2* CKO CMs than control CMs. *Fzd2* KD also increased YAP activity in NVCMs while *Fzd2* overexpression had the opposite effect. Together, these data suggest that FZD2 normally blocks CM proliferation by repressing YAP in these cells.

β-catenin binds YAP and enhances its induction of pro-proliferative genes in embryonic CMs suggesting that the high levels of β-catenin in *Fzd2* CKO CMs may activate YAP in these cells. We were thus surprised to find that β-catenin KD increased YAP protein and activity in NVCMs. These data and the lack of nuclear accumulation of β-catenin are inconsistent with β-catenin potentiating YAP-mediated transcription in *Fzd2* CKO CMs. Alternatively, YAP binds β-catenin in the destruction complex, raising the possibility that FZD2 promotes YAP degradation in the cytosol by increasing its affinity for β-catenin. However, this scenario is unlikely to be the sole mechanism for FZD2 function in regulating YAP as we found that FZD2 partially inhibited YAP in NVCMs after β-catenin KD. The effects of FZD2 and β-catenin on YAP activity thus appear to reflect at least partially independent processes.

YAP activation has been shown to improve cardiac outcomes in mice subject to MI^17–19^. The increased YAP activity in *Fzd2* CKO CMs may thus similarly improve cardiac function after ischemic injury. Consistent with this idea, *Fzd2* CKO mice had better EF and FS percentages as well as reduced fibrotic scaring than *Fzd2* HET mice after MI. *Fzd2* CKO CMs had increased cell cycle activity relative to controls after MI, suggesting that some of the damaged CMs are regenerated. However, cardiac function was improved in *Fzd2* CKO mice after only 3 days of recovery. Since it is unclear if new CMs could be produced fast enough to restore function within this timeframe, FZD2-deletion likely has both cardioprotective and reparative effects.

The AAR and extent of necrosis after MI were unaffected by FZD2-deletion, ruling out changes in the coronary vasculature that prevent the myocardium from becoming infarcted and *Fzd2* CKO CMs being insensitive to ischemia. Alternatively, changes in the inflammatory response may improve cardiac function and reduce scarring in *Fzd2* CKO hearts. FZD2-deletion may prevent CMs from releasing damage-associated molecular patterns that recruit neutrophils, which cause the secondary apoptosis of damaged CMs^71,72^. KD of WNT2/4 in mice has previously been shown to be cardioprotective against MI, and FZD2 was implicated in this process through activating NF-kB-dependent cardiac fibrosis using neonatal rat cardiac fibroblasts^73^, therefore *Fzd2* CKO CMs may also express paracrine factors that prevent cardiac fibroblasts from depositing collagen-rich scar tissue. Further studies will be needed to determine the relative contributions of cardioprotection and/or regeneration to the improved outcomes of *Fzd2* CKO mice after MI.

In summary, deleting *Fzd2* from the CMs of adult mice caused these normally post-mitotic cells to reenter the cell cycle. The pro-proliferative transcription factor YAP and its targets were enriched in CMs lacking FZD2 while FZD2 overexpression inhibited YAP activity in NVCMs. Although β-catenin levels were higher in FZD2-deficient CMs than controls, β-catenin localized to the membrane and did not activate TCF-dependent transcription in these cells. Moreover, β-catenin KD increased YAP activity in NVCMs and did not prevent FZD2 from inhibiting YAP in NVCMs. Furthermore, CM-specific FZD2 deletion dramatically improved heart function and reduced scarring in mice subject to MI without preventing the initial ischemic injury, suggesting that FZD2 inhibition may have therapeutic benefits for heart failure patients.

## Non-standard Abbreviations and Acronyms

AAR: Area at risk
BCL2L1: B cell leukemia/lymphoma 2
BrdU: 5-bromo-2’-deoxyuridine
CM: Cardiomyocyte
CKO: Conditional knockout
EF: Ejection fraction
FS: Fractional shortening
FZD2: Frizzled-2
GSK3αβ: Glycogen synthase kinase α and β
HW:TL: Heart weight to tibia length
I/R: Ischemia-reperfusion
KD: Knockdown
HET: Heterozygous
LATS1/2: Large tumor suppressors 1 and 2
MI: Myocardial infarction
MYCL: v-myc avian myelocytomatosis viral oncogene 1
MST1/2: Mammalian STE20-like protein kinases 1 and 2
NVCM: Neonatal ventricular cardiomyocyte
PBS: Phosphate buffered saline
PCM1: Pericentriole material 1
PFA: Paraformaldehyde
Q-PCR: Quantitative real timePCR
RPP: Rate pressure product
S.D.: Standard deviation
S.E.M.: Standard error of the mean
TAM: Tamoxifen
TCF: T-cell factor
TEAD: TEA-domain transcription factor
TTC: Triphenyl tetrazolium chloride
WGA: Wheat germ agglutinin
WT: Wild type
YAP: YES-associated protein

## SUPPLEMENTAL METHODS

### Isolation and culture of CMs and NVCMs

Adult mouse ventricular CMs were isolated as described^74^ with minor modifications. Briefly, hearts were removed and the aortas cannulated on a blunted 21G needle. The hearts were “hung” on a Langendorff apparatus and perfused with 1mg/mL collagenase II (Worthington Biochemical Corp, LS004176) at 37°C to dissociate CMs. Hearts were removed, the atria discarded and the ventricular tissue shredded with forceps before being passed through a 100 µm filter. CMs were separated by gravity sedimentation for 15 minutes at 37°C and the supernatant removed. The pelleted CMs were fixed in 2% paraformaldehyde or lysed in Trizol for RNA isolation. Fixed CMs were imaged using bright-field microscopy and CM areas were measured using Fiji ImageJ software. The numbers of nuclei were determined by manually counting DAPI-stained CMs using fluorescent microscopy. NVCMs were isolated from the hearts of neonatal mice as previously described^75^. NVCMs were plated in DMEM/F12 (Thermo Fisher, 11320) with 10% horse serum, 5% fetal bovine serum, antibiotic-antimycotic, Glutamax, and HEPES (Thermo Scientific, 16050, 16000, 15240, 35050 and 15630, respectively) on dishes coated with 25 µg/ml fibronectin (Thermo Fisher, 33016) in 0.1% gelatin solution (Millipore, ES-006-B).

### NVCM transfection and reporter assays

NVCMs were transfected with pools of siRNA against FZD2, β-catenin, or non-targeting control siRNA (Dharmacon, L-040443, L-040628 and D-0006-14, respectively) 24 hours after plating using Lipofectamine RNAiMAX (Thermo Fisher, 13778030) according to the manufacturer’s protocol. After recovering from siRNA transfection for 24 hours, NVCMs were transfected with the TEAD1-GAL4 reporter, which consists of pCMX-GAL4-Tead1 and pGL2-GAL4-UAS-Luc (AddGene, 33108 and 33020, respectively), or the 8x-TEAD-Luc reporter (AddGene, 34615) with Lipofectamine 2000 (Thermo Fisher, 11668027). NVCMs were also transfected with pRK5-*mFzd2* and pcDNA-Flag-*Yap1* (AddGene, 42254 and 18881, respectively) and pCMV-Tag2 (Agilent Technologies, 211172) for empty vector controls. All transfections included pCMV-SPORT-β-gal (Thermo Fisher, 10586014) to normalize for transfection efficiency. After 48 hours, NVCMs were lysed and analyzed with the Beta-Glo and Luciferase Assay Systems according to the manufacturer’s protocols (Promega Corp., E4720 and E1500, respectively).

### Q-PCR and western blotting

RNA was isolated from adult CMs and whole hearts with Trizol (Thermo Fisher, 15596026). Trizol was also used to isolate RNA from NVCMs 72 hours after siRNA transfection. qRT-PCR was performed with the ΔΔCT method as described^42^ using the primers listed in Supplemental Table 1. The Ct values of *Gapdh*, *Pol2ra*, and *Tbb* for each sample were averaged and used to normalize for loading. Protein was extracted from whole hearts by homogenizing tissue in RIPA supplemented with the HALT protease and phosphatase inhibitor cocktail (Thermo Fisher, 78440). Protein was obtained from NVCMs by scraping cells in RIPA with protease and phosphatase inhibitors 72 hours after siRNA transfection. Lysates were subject to western blotting as described^42^ with the antibodies listed in Supplemental Table 2. Band intensities were quantified using ImageJ 2.0 and normalized to the intensity of GAPDH.

### Histology and immunological staining

Hearts were flushed with 6 mg/l heparin in PBS to remove blood cells and prevent coagulation. Hearts were then perfused with 4% paraformaldehyde (PFA) and fixed overnight at 4°C before being prepared for histological processing as described^42^. Immunological staining was done as described^42,75^ using the primary antibodies in Supplemental Table 3. A biotin-conjugated secondary antibody and Alexa 488 Tyramide amplification kit (Thermo Fisher, B40932) were used to detect BrdU. Texas Red conjugated WGA (Thermo Fisher, W21405) was applied with secondary antibody. Slides were counterstained with 0.2 µg/ml DAPI in PBS and mounted in ProLong Gold (Thermo Fisher P36930). ImageJ 2.0 was used to quantify the cross-sectional areas of adult CMs and intensities of YAP staining in NVCMs. The collagen-rich scar tissue in post-MI samples was labeled with Masson’s trichrome (American Mastertech, KTMTR) and quantified with Adobe Photoshop. The numbers of BrdU+, Ki67+, and *TCF/LEF-GFP+* CMs were counted in images from an Olympus IX81 laser scanning confocal microscope.

### Echocardiography

To assess cardiac function, mice were anesthetized with isoflurane and subject to transthoracic echocardiography using a VisualSonics Vevo 2210. Readings were obtained with a 40 MHz transducer in M-mode along the short axis of the LV. Measurements were made by the staff of the Microsurgery and Echocardiography Core at the Aab Cardiovascular Research Institute, who were blinded to the genotype of the animals. To assess the sizes of the AAR and zone of necrosis, hearts were removed and suspended on a Langendorff apparatus 24 hours after MI. Hearts were perfused with PBS containing fluorescent microspheres (Thermo Scientific G0100) and Evans blue, cut into 2mm slices and incubated in 1% TTC (MP Biomedical Corp, 103126) for 15 minutes at 37°C before being fixed in 10% neutral buffered formalin (NBF).

### Statistics

Data is represented as the mean or median and interquartile range, where indicated, of n≥3 (unless otherwise indicated) independent biological replicates; error bars represent standard deviation (S.D.) or standard error of the mean (S.E.M.) where indicated. Student’s *t*-tests and one- or two-way ANOVA with Tukey’s multiple comparisons test were performed using SAS JMP and/or GraphPad Prism. Graphs were generated using GraphPad Prism.

---

**Supplemental Table 1:**
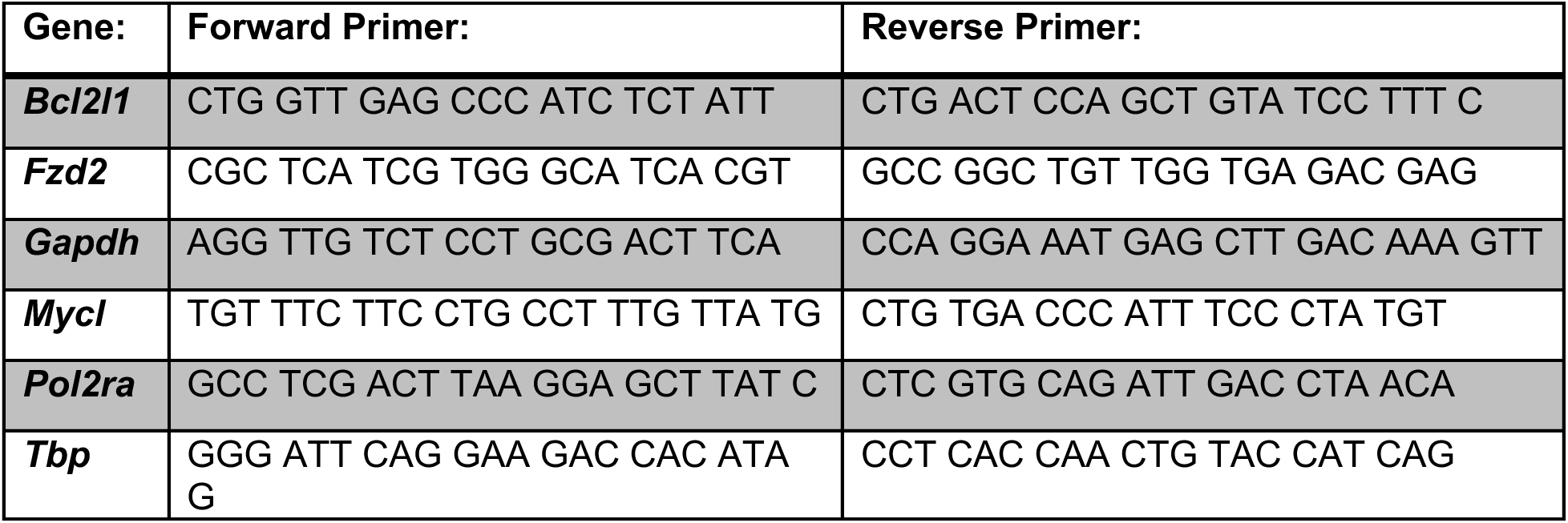
Primers used for quantitative real-time PCR (Q-PCR) listed 5’ to 3’.

**Supplemental Table 2:** Antibodies used with manufacturer’s names and catalog numbers.

|  |  |  |
| --- | --- | --- |
| Antibody: | Manufacturer: | Catalog #: |
| **Aurora B kinase** | Abcam | ab2254 |
| **BrdU** | Novus | NB500-169 |
| **α-Cardiac Actin (ACTC1)** | Thermo Fisher | MA1-10046 |
| **β-catenin (non-phosphorylated)** | Cell Signaling | 4270S |
| **β-catenin (total)** | BD Transduction Laboratories | 610154 |
| **YAP (IHC)** | Santa Cruz | SC-101199 |
| **YAP/TAZ (western blots)** | Cell Signaling | 8418P |
| **YAP (phospho-S127)** | Cell Signaling | 13008 |
| **YAP (phospho-S397)** | Cell Signaling | 13619P |
| **Ki67** | eBioscience | 14-5698-80 |
| **FZD2** | Sigma-Aldrich | SAB2501520 |
| **Desmin** | Invitrogen | PA1-27003 |
| **GAPDH** | Cell Signaling | 2118S |
| **GFP** | Thermo Fisher | MA5-15256 |
| **Cardiac Troponin T (TNNT2)** | Abcam | Ab8295 |
| **Desmoplakin** | Proteintech | 25318-1-AP |

**Supplemental Table 3:** Cardiac function in *Fzd2* HET and *Fzd2* CKO mice before (baseline) and after myocardial infarction as assessed by serial transverse echocardiography. Chart shows the mean ± S.E.M. for each parameter. Asterisks indicate p ≤ 0.05 on two-way ANOVAs with Tukey’s multiple comparison tests.

| Parameter | Baseline | MI D3 | MI D7 | MI D14 | MI D28 |
| --- | --- | --- | --- | --- | --- |
| **Heart rate (beats per minute)** |  |  |  |  |  |
| HET | 537.16 ± 21.69 | 534.64 ± 27.91 | 560.89 ± 36.93 | 518.55 ± 9.80 | 556.03 ± 14.29 |
| CKO | 534.29 ± 31.56 | 563.37 ± 31.13 | 531.59 ± 44.37 | 562.69 ± 35.02 | 534.43 ± 36.17 |
| **LV Ejection fraction (%)** |  | * | * | * | * |
| HET | 73.87 ± 15.38 | 28.69 ± 19.91 | 32.49 ± 16.90 | 30.09 ± 11.40 | 18.48 ± 8.49 |
| CKO | 68.08 ± 11.40 | 65.72 ± 19.04 | 61.49 ± 17.66 | 61.95 ± 15.66 | 55.69 ± 18.53 |
| **LV Fractional shortening (%)** |  | * | * | * | * |
| HET | 43.89 ± 13.11 | 13.85 ± 10.33 | 16.06 ± 9.49 | 14.56 ± 5.79 | 8.63 ± 4.16 |
| CKO | 40.35 ± 9.69 | 42.10 ± 3.49 | 38.02 ± 6.19 | 36.22 ± 6.90 | 32.37 ± 7.76 |
| **LV internal diameter, systole (mm)** |  | * | * | * | * |
| HET | 2.17 ± 0.67 | 3.99 ± 1.05 | 4.52 ± 1.10 | 4.76 ± 0.49 | 5.70 ± 0.62 |
| CKO | 2.47 ± 0.65 | 2.27 ± 0.37 | 2.43 ± 0.55 | 2.72 ± 0.36 | 2.72 ± 0.69 |
| **LV internal diameter, diastole (mm)** |  |  | * | * | * |
| HET | 3.58 ± 0.47 | 4.63 ± 0.77 | 5.26 ± 0.76 | 5.53 ± 0.44 | 6.19 ± 0.42 |
| CKO | 3.70 ± 0.65 | 3.87 ± 0.40 | 3.93 ± 0.48 | 4.26 ± 0.34 | 4.01 ± 0.57 |
| **LV anterior wall diameter, systole (mm)** |  |  |  | * | * |
| HET | 1.67 ± 0.20 | 1.16 ± 0.62 | 1.23 ± 0.53 | 0.94 ± 0.40 | 0.57 ± 0.17 |
| CKO | 1.69 ± 0.20 | 1.72 ± 0.37 | 1.48 ± 0.19 | 1.72 ± 0.29 | 1.63 ± 0.36 |
| **LV anterior wall diameter, diastole (mm)** |  |  |  |  | * |
| HET | 1.18 ± 0.19 | 0.94 ± 0.40 | 1.11 ± 0.34 | 0.82 ± 0.27 | 0.58 ± 0.21 |
| CKO | 1.24 ± 0.20 | 1.23 ± 0.21 | 0.99 ± 0.09 | 1.19 ± 0.23 | 1.14 ± 0.23 |
| **LV posterior wall diameter, systole (mm)** |  |  |  |  |  |
| HET | 1.57 ± 0.23 | 1.36 ± 0.41 | 1.45 ± 0.33 | 1.20 ± 0.39 | 1.15 ± 0.49 |
| CKO | 1.61 ± 0.13 | 1.66 ± 0.28 | 1.68 ± 0.21 | 1.55 ± 0.41 | 1.47 ± 0.41 |
| **LV posterior wall diameter, diastole (mm)** |  |  |  |  |  |
| HET | 1.08 ± 0.05 | 1.13 ± 0.26 | 1.17 ± 0.31 | 1.00 ± 0.21 | 0.90 ± 0.36 |
| CKO | 1.25 ± 0.17 | 1.11 ± 0.24 | 1.19 ± 0.11 | 1.07 ± 0.30 | 1.14 ± 0.25 |
| **Volume, systole (μL)** |  | * | * | * | * |
| HET | 17.03 ± 13.12 | 77.74 ± 43.80 | 100.76 ± 49.02 | 107.79 ± 27.49 | 160.95 ± 37.42 |
| CKO | 22.54 ± 18.21 | 17.93 ± 7.13 | 23.62 ± 12.19 | 28.65 ± 12.21 | 32.08 ± 17.08 |
| **Volume, diastole (μL)** |  |  | * | * | * |
| HET | 61.17 ± 15.72 | 101.43 ± 36.79 | 141.04 ± 42.07 | 153.55 ± 26.63 | 195.60 ± 27.35 |
| CKO | 67.91 ± 24.64 | 65.76 ± 15.29 | 71.95 ± 21.46 | 80.84 ± 12.60 | 74.47 ± 25.34 |

**Supplemental Figure 1.**
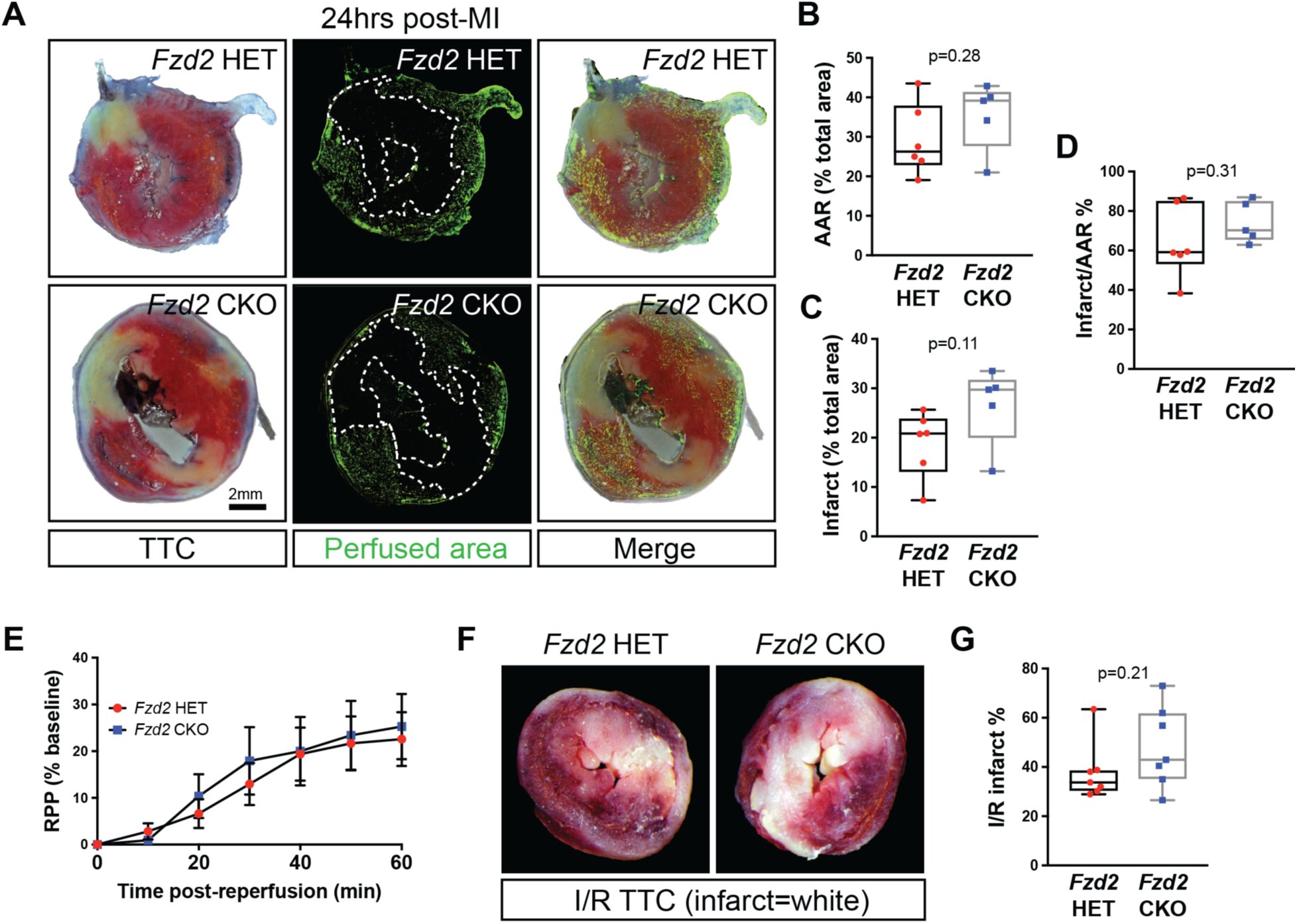
FZD2-deletion does not prevent acute ischemic injury after infarction. **A.** *Fzd2* HET (n = 6) and *Fzd2* CKO (n = 5) mice were subject to experimental MI. After 24 hours, hearts were cannulated and perfused with 1 µm fluorescent micro-spheres and Evan’s blue to determine area at risk (AAR). TTC staining was used to assess the extent of necrosis. **B.** Quantification of AAR. **C.** Infarct size. **D.** Infarct/AAR ratio (**D**). **E-G.** Hearts from *Fzd2* HET and *Fzd2* CKO mice were cannulated and perfused with oxygenated buffer for 25 minutes to equilibrate. Flow was then turned off to induce ischemia for 20 minutes before being restored for 1 hour to simulate reperfusion. **E.** Percent change in rate pressure product (RPP) over time starting at reperfusion. **F.** TTC staining from hearts after I/R. **G.** Infarct size quantification from TTC staining after I/R. All graphs show the mean ± S.E.M, p-values are as indicated by Student’s t-tests.

